# Identification of hippocampal area CA2 in hamster and vole brain

**DOI:** 10.1101/2024.02.12.579957

**Authors:** Preston Nicole Siegler, Emma K. Shaughnessy, Brian Horman, Tia T. Vierling, Darron H. King, Heather B. Patisaul, Kim L. Huhman, Georgia M. Alexander, Serena M. Dudek

## Abstract

Prairie voles (*Microtus ochrogaster*) and Syrian, or golden, hamsters (*Mesocricetus auratus*) are closely related to mice (*Mus musculus*) and rats (*Rattus norvegicus*, for example) and are commonly used in studies of social behavior including social interaction, social memory, and aggression. The CA2 region of the hippocampus is known to play a key role in social memory and aggression in mice and responds to social stimuli in rats, likely owing to its high expression of oxytocin and vasopressin 1b receptors. However, CA2 has yet to be identified and characterized in hamsters or voles. In this study, we sought to determine whether CA2 could be identified molecularly in vole and hamster. To do this, we used immunofluorescence with primary antibodies raised against known molecular markers of CA2 in mice and rats to stain hippocampal sections from voles and hamsters in parallel with those from mice. Here, we report that, like in mouse and rat, staining for many CA2 proteins in vole and hamster hippocampus reveals a population of neurons that express regulator of G protein signaling 14 (RGS14), Purkinje cell protein 4 (PCP4) and striatal-enriched protein tyrosine phosphatase (STEP), which together delineate the borders with CA3 and CA1. These cells were located at the distal end of the mossy fiber projections, marked by the presence of Zinc Transporter 3 (ZnT-3) and calbindin in all three species. In addition to staining the mossy fibers, calbindin also labeled a layer of CA1 pyramidal cells in mouse and hamster but not in vole. However, Wolframin ER transmembrane glycoprotein (WFS1) immunofluorescence, which marks all CA1 neurons, was present in all three species and abutted the distal end of CA2, marked by RGS14 immunofluorescence. Staining for two stress hormone receptors—the glucocorticoid (GR) and mineralocorticoid (MR) receptors—was also similar in all three species, with GR staining found primarily in CA1 and MR staining enriched in CA2. Interestingly, although perineuronal nets (PNNs) are known to surround CA2 cells in mouse and rat, we found that staining for PNNs differed across species in that both CA2 and CA3 showed staining in voles and primarily CA3 in hamsters with only some neurons in proximal CA2 showing staining. These results demonstrate that, like in mouse, CA2 in voles and hamsters can be molecularly distinguished from neighboring CA1 and CA3 areas, but PNN staining is less useful for identifying CA2 in the latter two species. These findings reveal commonalities across species in molecular profile of CA2, which will facilitate future studies of CA2 in these species. Yet to be determined is how differences in PNNs might relate to differences in social behavior across species.

## Introduction

Within the hippocampus proper, the *Cornu Ammonis* 2 (CA2) region has historically been defined as the CA region between the end of the mossy fiber inputs from dentate gyrus granule cells and area CA1 (Lorente de Nò, 1934). Like in CA3, CA2 contains pyramidal neurons with somas larger than those of neighboring CA1 neurons, but unlike in CA3, the pyramidal cell dendrites of CA2 cells largely lack the characteristic thorny excrescences found associated with mossy fiber inputs on CA3 neurons (Lorente de Nò, 1934; Ramón y Cajal, 1911). Although the CA subregions were named by Lorente de Nó, the initial definition of CA2 has since been at odds with the more recent molecular definition, which more closely matches the CA2 neurons in drawings by Ramón y Cajal, showing cells lacking the complex spines that also receive input from the mossy fibers (Kohara et al., 2014; Ramón y Cajal, 1911). In current studies, which have used primarily mice and rats, molecular markers such as regulator of G protein signaling 14 (RGS14), striatal-enriched protein tyrosine phosphatase (STEP) and Purkinje cell protein 4 (PCP4) are immunostained to demarcate the borders of CA2 with CA1 and CA3 and can also be used to examine CA2 projections in isolation (Kohara et al., 2014; Radzicki et al., 2023).

Neurons in CA2 have a variety of properties that make them unique in comparison to those in other areas of the hippocampus. Area CA2 is rich in proteins that suppress long term potentiation (LTP), including RGS14, STEP, and likely PCP4 (Boulanger et al., 1995; Lee et al., 2010; Lein et al., 2005; Pelkey et al., 2002; Simons et al., 2009). As such, LTP is nearly nonexistent at excitatory synapses in CA2 *stratum radiatum* (SR; although robust LTP is seen in *stratum lacunosum-moleculare*)(Chevaleyre & Siegelbaum, 2010; Zhao et al., 2007). However, synaptic potentiation at SR synapses does occur upon pharmacological activation of vasopressin or oxytocin receptors, which are both enriched in mouse and rat CA2 (Lee, Caldwell, Macbeth, Tolu, et al., 2008; Lee, Caldwell, Macbeth, & Young, 2008; Pagani et al., 2015; Tirko et al., 2018; Wersinger et al., 2002; Young et al., 2006). These findings, alone, suggest that CA2 could play an important role in social learning and behavior. Indeed, social memory and aggression are impaired in mice lacking vasopressin 1B receptors, which are enriched in CA2 (DeVito et al., 2009; Wersinger et al., 2002). Similarly, CA2 silencing impairs social memory and aggression (Hitti & Siegelbaum, 2014; Leroy et al., 2018).

Although mice are commonly used in neuroscience research because of their ease of genetic manipulation, prairie voles (*Microtus ochrogaster*) and Syrian hamsters (*Mesocricetus auratus*) are also used in behavioral studies of social learning and memory, pair bonding, and aggression due to their specific behavioral characteristics. For example, prairie voles exhibit monogamous pair bonding and alloparenting behavior, so they are valuable model organisms of prosocial behaviors (Gillera et al., 2022; Stetzik et al., 2018). In contrast, both male and female Syrian hamsters are naturally antagonistic, which makes them ideal for study of social dominance and aggression (Cooper et al., 2009; Solomon et al., 2009; Taylor et al., 2022).

Despite the common use of prairie voles and Syrian hamsters in social behavior research and the now-appreciated role of CA2 in social behavior in mice and rats, it remains unknown whether prairie voles and Syrian hamsters possess a hippocampal structure homologous to mouse CA2. Recognition and characterization of a vole and hamster area CA2 would permit better generalization of findings from mouse to other species, future study of the neuronal underpinnings of social behaviors in voles and hamsters, and a better understanding of the role of CA2 in prosocial and aggressive behaviors. Therefore, in this study, we sought to characterize the molecular profile of vole and hamster CA2, its anatomical position within the CA subfields, and other anatomical features as they relate to observations in mouse CA2.

## Materials and Methods

### Animal Care and Handling

All animals were used in accordance with the standards set by the Animal Welfare Act and the U.S. Department of Health and Human Services “Guide for the Care and use of Laboratory Animals”. Care and procedures were approved by the respective Institutional Animal Care and Use Committees at North Carolina State University (NCSU; voles), Georgia State University (GSU; hamsters), or the National Institute of Environmental Health Sciences (NIEHS; mice). Four male and four female unbonded, untreated wild type voles were cared for as in Patisaul, et al. (Patisaul & Adewale, 2009; Patisaul et al., 2013). Food and water were provided ad libitum and animals were group housed in animal rooms with 12 hour light/dark cycles in the Assessment and Accreditation of Laboratory Animal Care approved Biological Resource Facility at NCSU.

Three male and four female untreated wildtype Syrian hamsters *(M. auratus)* were cared for as published in Partrick et al., 2021 (Partrick et al., 2021). Syrian hamsters were purchased from Charles River Laboratories and single housed in a 14-hour light/10-h dark cycle, as is common in hamsters to maintain gonadal patency, with ad libitum food and water.

Voles and hamsters were naive to any treatment at the time of perfusion. At their respective institutions, 4-month-old hamsters and 4-month-old voles were perfused using 4% paraformaldehyde (PFA) via cardiac puncture. Brains were removed and post-fixed for at least 24 hours before transporting to NIEHS in glycerol-based cryoprotectant or 4% PFA for voles and hamsters respectively. Brains were then cut coronally at 40 µM using a vibratome and stored in PBS with sodium azide.

Male and female wild type mice were group housed at NIEHS under a 12:12 light/dark cycle with ad libitum access to food and water. Mice were naive to any treatment at the time of perfusion. Animals used in histology were euthanized with Fatal Plus (sodium pentobarbital, 50 mg/mL; >100 mg/kg) and immediately transcardially perfused with 4% PFA, and brains remained in solution until sectioning. Brains were then sectioned coronally with a vibratome at 40 μm and stored in PBS with sodium azide.

### Immunostaining and Imaging

Immunofluorescence (IF) staining was performed as published in Alexander, et al. 2018 (Alexander et al., 2018). Brain sections were washed once for 10 minutes in PBS, and for antigen retrieval (used in all stains but the *Wisteria Floribunda* agglutinin, WFA, and aggrecan, ACAN, staining), sections were placed in 1 mL microfuge tubes along with 0.75 ml of deionized water. These tubes were then boiled in deionized water for 2 min and sections subsequently blocked for 1 hour in 3% normal goat serum (NGS)/0.1% triton x100 PBS (PBST). Sections were incubated in primary antibodies as listed in Table 1. Antibodies were diluted in either a 3% or 5% NGS blocking solution, and sections were incubated on a rocker in a 4℃ cold room for a minimum of 12 hours and a maximum of 24 hours. After several rinses in PBST, sections were incubated in secondary antibodies as listed in Table 2 for 3 hours with antibodies diluted as in primary, in either a 3% or 5% NGS blocking solution. Sections were then washed in PBST and mounted using either Vectashield Antifade Mounting Medium with DAPI (coverslip sealed with clear instant dry nail polish) or with Vectashield HardSet Antifade Mounting Medium with DAPI (Vector Laboratories). Imaging was performed using a Zeiss Axio Observer microscope and the resulting photos pseudo-colored for clarity using ImageJ software obtained from the National Institutes of Health (Schneider et al., 2012). The sagittal and horizontal RGS14 and PCP4 staining, the 40x oil images of the WFA and ACAN staining, and the receptor staining were imaged on a Zeiss 980 Confocal microscope. Obtained images were processed as listed above. In most cases brightness was optimized for illustrative purposes and were not necessarily matched between species unless noted.

**Table 1:** List of primary antibodies utilized in this study with company, host species, reference and lot numbers, clonality, immunogen information, Research Resource Identifiers, and dilution used for staining.

| NAME | HOST SPECIES | COMPANY | REF/lot | CAT # | CLONALITY or TYPE | ANTIGENS | RRID | DILUTION |
| --- | --- | --- | --- | --- | --- | --- | --- | --- |
| Aggrecan | Rabbit | Millipore | 3800006 | AB1031 | Polyclonal | GST fusion protein containing mouse aggrecan aa 1177-1326 | AB_90460 | 1:500 |
| Calbindin | Rabbit | Swant | 38 | CB-38a | Polyclonal | Recombinant rat calbindin D-28k (CB) | AB_10000340 | 1:500 |
| GR | Mouse | Santa Cruz Biotechnology | D2923 | sc-393232 | Monoclonal | GR (G-5) – human | AB_2687823 | 1:250 |
| MR | Mouse | Millipore | 3940936 | MABS496 | Monoclonal | Conjugated linear peptide corresponding to rat Mineralocorticoid Receptor, clone 6G1. | AB_2811270 | 1:100 |
| PCP4 | Guinea Pig | Synaptic Systems | 1-1 | 480 004 | Polyclonal | Full length human recombinant PCP4 | AB_2927387 | 1:500 |
| PCP4 | Rabbit | Santa Cruz Biotechnology |  | sc-98549 | Polyclonal | Peptide mapping to the C-terminus of human PCP4 (C-15) | AB_2251885 | 1:500 |
| RGS14 | Mouse | Neuromab | clone n133/21 | N133/35 | Monoclonal | Fusion protein of full-length rat RGS14, epitope mapped to aa 444-490 | AB_2877596 | 1:500 |
| STEP | Mouse | Cell signaling | 2 | 4396 | Monoclonal | Synthetic peptide corresponding to the amino terminus of rat STEP46 protein | AB_1904101 | 1:500 |
| WFA | N/A | Vector | b-1355 | Zg0218 | Lectin | Species N/A | AB_2336874 | 1:500 |
| WFS1 | Rabbit | Proteintech | 00043335 | 11558-1-AP | Polyclonal | Human WFS1 fusion protein Ag2114 | AB_2216046 | 1:500 |
| ZnT-3 | Guinea Pig | Synaptic Systems | 197004 | ZnT-3 | Polyclonal | Recombinant protein corresponding to aa 2-75 from mouse ZnT3 | AB_2189667 | 1:500 |

**Table 2:** Table 2: List of secondary antibodies use in this study with company, animal, reference and lot numbers, and dilution used for staining.

| NAME | ANIMAL<br>recognized | COMPANY | REF/lot | CAT | DILUTION |
| --- | --- | --- | --- | --- | --- |
| Alexa Fluor 488 | Rabbit | Invitrogen | 2156517 | A11034 | 1:500 |
| Alexa Fluor 568 | Mouse | Invitrogen | 2300933 | A11031 | 1:500 |
| Alexa Fluor 568 | Mouse | Invitrogen | 2332536 | A11004 | 1:500 |
| Alexa Fluor γ1 488 | Mouse | Invitrogen | 2339820 | A21121 | 1:500 |
| Alexa Fluor γ2a<br>555 | Mouse | Invitrogen | 2335727 | A21137 | 1:500 |
| Alexa Fluor 488 | Guinea Pig | Invitrogen | 982288 | A11073 | 1:500 |
| Alexa Fluor 633 | N/A, Biotin | Invitrogen | 1728204 | S21375 | 1:500 |

### Antibodies Used

Primary and secondary antibodies in this study are listed in Tables 1 and 2, respectively, with their Research Resource Identifier (RRID) numbers, immunogen information from the manufacturers’ product sheets, catalogue and lot numbers, and the dilutions used. Of the antibodies used, RGS14, ZnT-3, and calbindin were verified with knock-out mouse experiments (Celio et al., 1990; Girard et al., 2015; Lee et al., 2010; Li et al., 2017) and others, such as ACAN, the glucocorticoid receptor (GR), the mineralocorticoid receptor (MR), PCP4, STEP, and WFS1 were validated via western blot, per information from the manufacturers’ datasheets and as previously reported (Erhardt et al., 2000; Gomez-Sanchez et al., 2006; Paul et al., 2000; Sasse et al., 2015; Yi et al., 2012). ZnT-3 and RGS14 were also validated with western blots per the manufacturers’ datasheets.

## Results

We used IF staining to visualize cellular populations in this study. Many previous studies studying brains from animals such as cats, hedgehogs, and guinea pigs, have utilized chromogenic immunohistochemistry (IHC) techniques such as diaminobenzidine (DAB) staining (Hirama et al., 1997; Kunzle & Radtke-Schuller, 2001; Rami et al., 1987; Zilli et al., 2022), which, although useful in identifying the cellular density of a region, does not provide the spatial or molecular specificity afforded by multiplexing with combinations of immunofluorophores. We therefore took advantage of IF staining using known markers of mouse CA2. We note that although we observed no obvious difference in staining between tissue from males and females, we made no attempt here to test whether there were sex differences.

### RGS14 immunostaining demarcates hippocampal area CA2 in mouse, vole, and hamster

Mouse, vole, and hamster brain sections were immunostained for RGS14, a commonly used marker of CA2 neurons. RGS14-positive cell populations were found in all animals and appeared in multiple planes of the hippocampus from anterior to posterior (Fig.1). Antibodies raised against RGS14 were used as the main CA2 marker for this study, as it is well documented as a reliable CA2 marker in mice (Lee et al., 2010; Radzicki et al., 2023). Tissue from all three species examined here showed consistent RGS14-labeled cells in an anatomical area similar to that defined as hippocampal area CA2 in mice in that the CA1/CA2 border is well defined, and the CA2/CA3 border somewhat less so, most evident in the hamster (Fig. 1) (Radzicki et al., 2023). Staining for DAPI, which binds to nuclear DNA, was used throughout the study to visualize cellular density and dispersion across hippocampal subfields. Note that mouse CA2 and CA3 are typically only lightly stained with DAPI. The overall size of these hippocampus sections, as well as the area of the associated RGS14-positive cellular populations, differed across the three species, with hamster having the largest hippocampus and thus CA2, followed by vole, and then mouse (Fig. 1), which is consistent with the differences in overall brain sizes. Although Figure 1 serves to contextualize this relative size difference, higher magnification views in subsequent figures allow closer examination of the RGS14-positive cell population.

**Figure 1:**
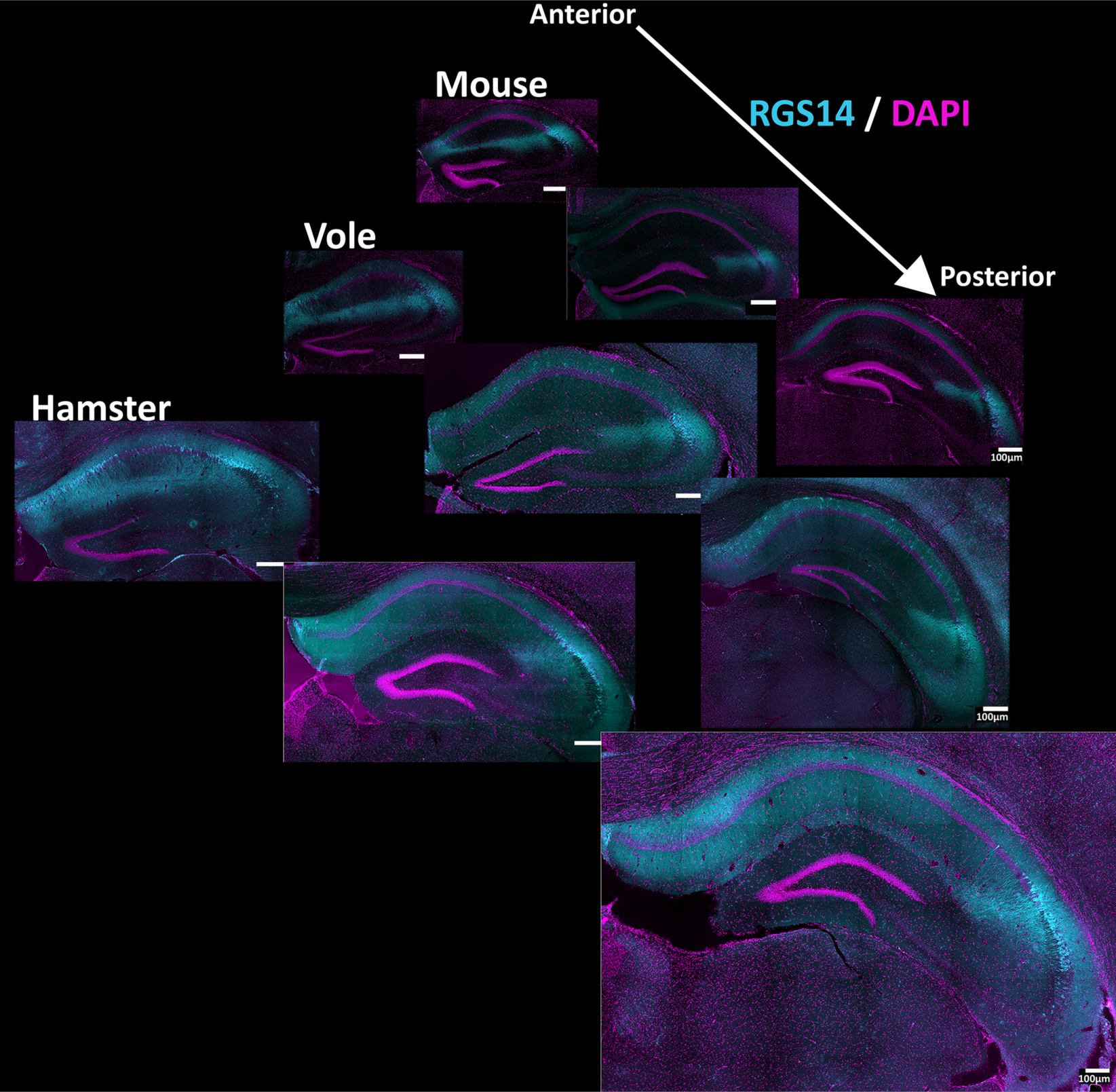
Anterior to posterior view of hippocampal area CA2 In hamster, vole, and mouse. RGS14 positive cell populations demarcate hippocampal area CA2 in hamster, vole, and mouse. Coronal hippocampal sections stained with RGS14 (cyan), a commonly used marker of hippocampal area CA2, indicate a cellular population inferred to be area CA2. DAPI (magenta) shows cellular density. Anterior, medial, and posterior example sections are shown for each species. Sections are scale-matched to show their relative sizes, with all scale bars set at 100 μm.

### Mossy fiber and CA1 visualization provide anatomical context for area CA2 in mouse, vole, and hamster

Molecularly defined CA2 neurons in mice are anatomically positioned at the distal end of the mossy fibers that pass through CA3 and receive functional input from the dentate granule neurons (Kohara et al., 2014). The tail (distal) end of CA2, just beyond the end of the mossy fibers, borders CA1. To identify the anatomical positioning of RGS14-defined CA2 relative to these landmarks in vole and hamster, we labeled the mossy fibers with ZnT-3 or calbindin antibodies, and labeled CA1 with an antibody recognizing WFS1. Considering that calbindin has also been shown to label the pyramidal cell layer of mouse CA1 closest to the SR (superficial), it was also used here alongside WFS1 to demarcate CA1 (Kohara et al., 2014). We found that ZnT-3 and calbindin both labelled in the mossy fiber projections from dentate gyrus in all three species (Fig. 2a-b) and RGS14-labeled neurons were positioned at the tail end of the mossy fibers. In addition, we found that the WFS1 antibody stained CA1 neurons in all species, and these cells bordered the distal end of RGS14-defined CA2. Therefore, as in mouse, CA2 neurons in vole and hamster are positioned between the tail end of the mossy fibers and CA1, supporting the conclusion that this population of neurons can be defined as CA2 in vole and hamster. We found that antibodies raised against WFS1 (Mody et al., 1987; Takeda et al., 2001) were most effective in labeling area CA1 in mouse and the weakest in hamster. Similarly, antibodies raised against ZnT-3 were most effective in labeling the mossy fiber system in mouse and weakest in hamster.

**Figure 2:**
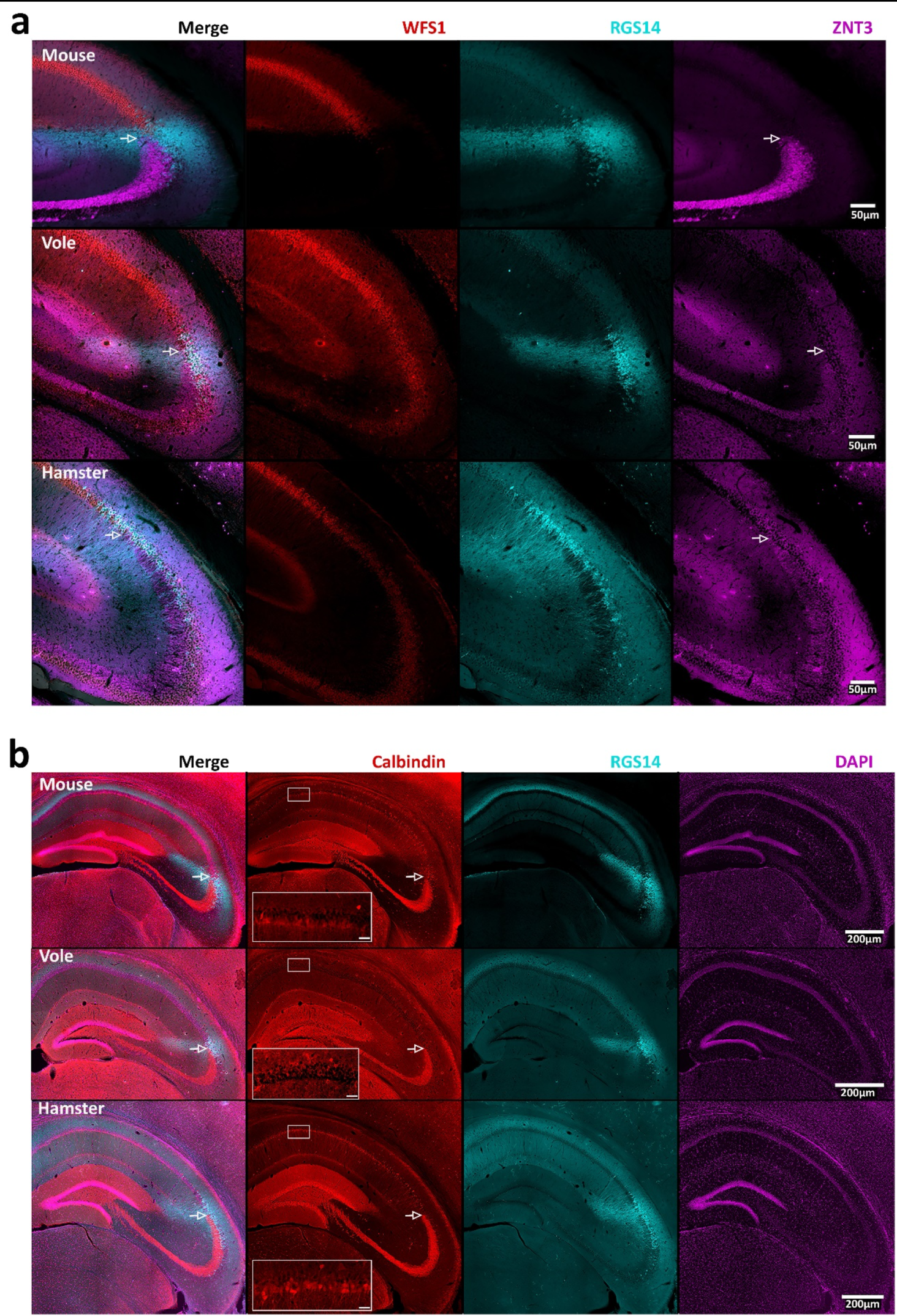
Visualization of hippocampal area CA2 in relation to mossy fibers and area CA1 pyramidal cells. (a) Images of coronal sections showing neurons in CA1 stained with WFS1 (red), with staining for RGS14 (cyan), indicating area CA2 and ZnT-3 (magenta) staining the mossy fibers. The arrows indicate the presumed ending of the mossy fibers. (b) Images showing mossy fibers labeled with an antibody against calbindin (red), which also label CA1 pyramidal cells. As in (a), arrows indicate their presumed end of the mossy fibers and RGS14 was used to visualize a cellular population inferred to be area CA2 (cyan). Cellular density was visualized using DAPI (magenta). The leftmost column (a, b) shows merged images. A 20x view of calbindin positive cells, or lack thereof, in CA1 is shown as an inset with scale bars set at 25 µm. Other scale bars are as indicated.

To further compare the location of CA2 neurons in the different species with respect to CA1, we used antibodies raised against calbindin, which, like WFS1, has been used to label CA1 pyramidal neurons in a variety of species (Mody et al., 1987; Takeda et al., 2001). Interestingly, we observed strong staining of the mossy fibers in all three species, but calbindin-positive cells in area CA1 varied; we saw strong staining of in mouse, varied staining in hamster, but no staining in CA1 in vole. A layer of calbindin-positive cells can be seen adjacent to a layer of calbindin-negative cells in both mouse and hamster (Fig. 2). Thus, although the calbindin antibody remains a good marker for the mossy fibers in the species tested here, it is less useful for marking the CA1/CA2 border in some species.

### RGS14 colocalizes with other common area CA2 markers

Although RGS14 is a robust marker of area CA2 in all species examined (Squires et al., 2018; Lee et al., 2010), we thought it could be beneficial to determine if this protein colocalizes with others that are commonly used as CA2 markers (Boulanger et al., 1995; Lee et al., 2010; Lein et al., 2005; Radzicki et al., 2023). We therefore co-stained sections with antibodies against RGS14 and PCP4 to probe for overlap in CA2. We found that staining for PCP4 overlapped with RGS14 staining in mouse, consistent with previous studies, and this was also the case in sections from voles and hamsters (Fig. 3). Despite an area of reliable colocalization of PCP4 and RGS14 staining in mice, prior research has also noted an incomplete overlap on the edges of the double-labelled zone, with scattered cells at the border with CA1 staining only for PCP4 and cells at the border with CA3 staining only for RGS14 (Radzicki et al., 2023). Similar staining patterns are visible in vole and hamster CA2 (Fig. 3). In addition, we found that both RGS14 and PCP4 were differentially expressed in dorsal and ventral CA2 in all three species (Fig. 4), consistent with previous observations in mouse (Bienkowski, 2023). Specifically, RGS14 appeared to be more highly expressed in dorsal CA2 than in ventral CA2, whereas by comparison, staining for PCP4 appeared more equal in both dorsal and ventral CA2, albeit still greater in dorsal CA2. This suggests that, like RGS14, PCP4 can be used as a marker of CA2 in mouse, vole, and hamster. Furthermore, staining for PCP4 more clearly identifies CA2 than RGS14 staining in the ventral hippocampus. Staining with another common CA2 marker, STEP (Supplemental Fig. 1), similarly showed expression that colocalized with RGS14 in dorsal CA2. As with other antibodies tested, staining for STEP was much more robust in mouse than in either vole or hamster.

**Figure 3:**
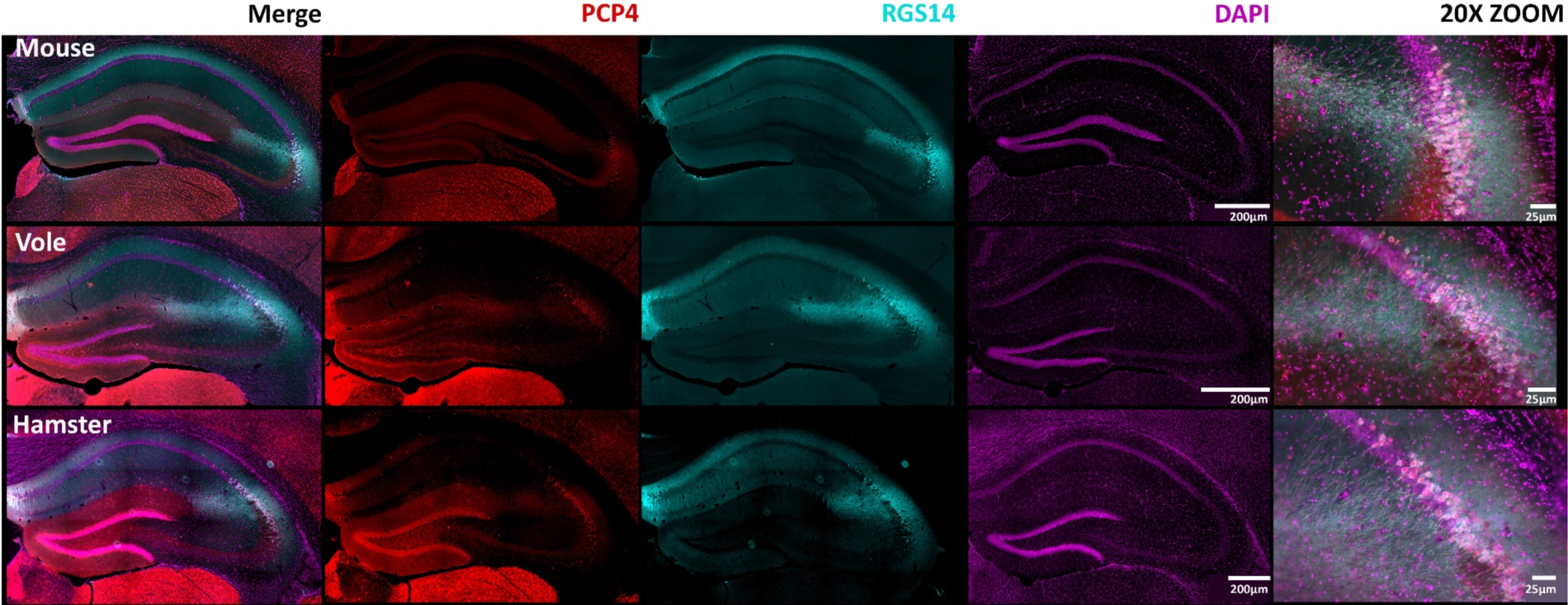
RGS14 colocalizes with another putative area CA2 marker-PCP4. Coronal sections of mouse, vole, and hamster hippocampus are shown. Purkinje cell protein 4 (PCP4; red) is present in all three species and colocalizes with RGS14 (cyan), demarcating area CA2, and DAPI (magenta) showing cellular density. Images of CA2 at 20x zoom are shown on the right. Scale bars are as indicated.

**Figure 4:**
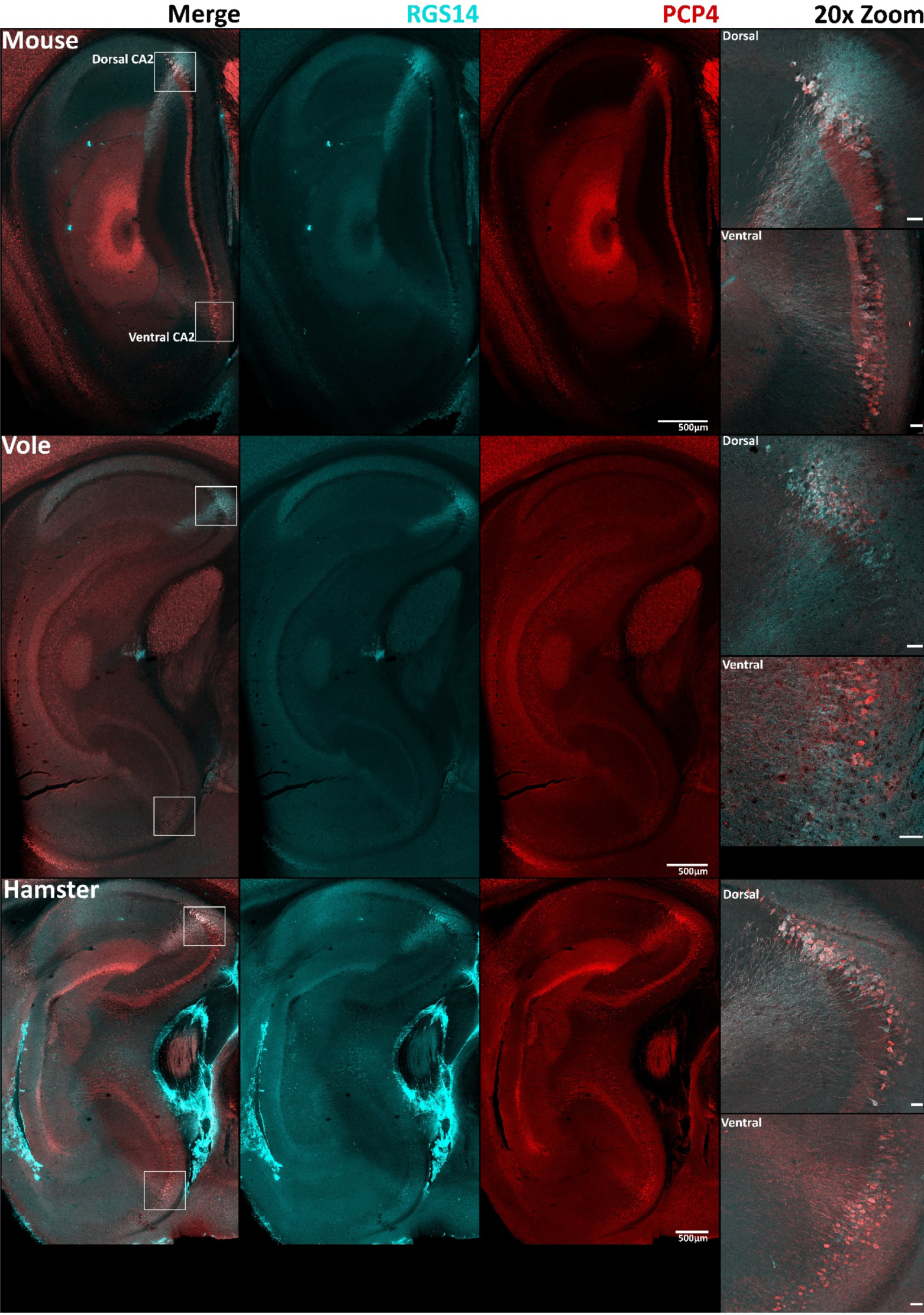
RGS14 and PCP4 are differentially expressed in dorsal and ventral CA2 in mouse, vole, and hamster. Lateral sagittal sections are shown to visualize both dorsal and ventral hippocampus. Both PCP4 (red) and RGS14 (cyan) are present in dorsal and ventral CA2 of all three species, but RGS14 appears much more intense in dorsal CA2 than in ventral CA2. The leftmost column shows merged images. A 20x zoom of both dorsal and ventral CA2 is shown on the right with scale bars set at 50 µm.

### Perineuronal net staining does not serve as a marker of CA2 in vole or hamster

Perineuronal nets (PNNs) are extracellular matrix structures that typically envelop inhibitory parvalbumin-expressing interneurons in the brain and spinal cord (Balmer et al., 2009; Hartig et al., 1992). In mouse, PNNs also surround CA2 pyramidal neurons (Carstens et al., 2016). Two common methods of staining PNNs in mouse tissue are with the lectin *Wisteria Floribunda* agglutinin (WFA), which directly labels PNNs (Bruckner et al., 2003; Celio, 1993; Celio & Chiquet-Ehrismann, 1993; Nadanaka et al., 2020; Radzicki et al., 2023), and antibodies raised against a primary proteoglycan component of PNNs, aggrecan (ACAN) (Carstens et al., 2016; Radzicki et al., 2023). We stained mouse, vole, and hamster sections with both WFA and an antibody against ACAN to label PNNs and co-stained tissue with RGS14. In all three species, we observed both ACAN and WFA staining surrounding presumptive parvalbumin-expressing interneurons scattered throughout the hippocampus. However, staining among pyramidal cells showed striking differences across species. Whereas mouse staining showed the expected colocalization of PNN markers with RGS14 within CA2, tissue from both vole and hamster showed PNN stain primarily surrounding CA3 pyramidal neurons, with only some RGS14 positive neurons staining evident in the area closest to CA3, likely representing the proximal portion of CA2 (Fig 5). These findings highlight some of the likely molecular differences between different rodent species. The functional consequences of these differences are unknown.

**Supplemental Figure 1:**
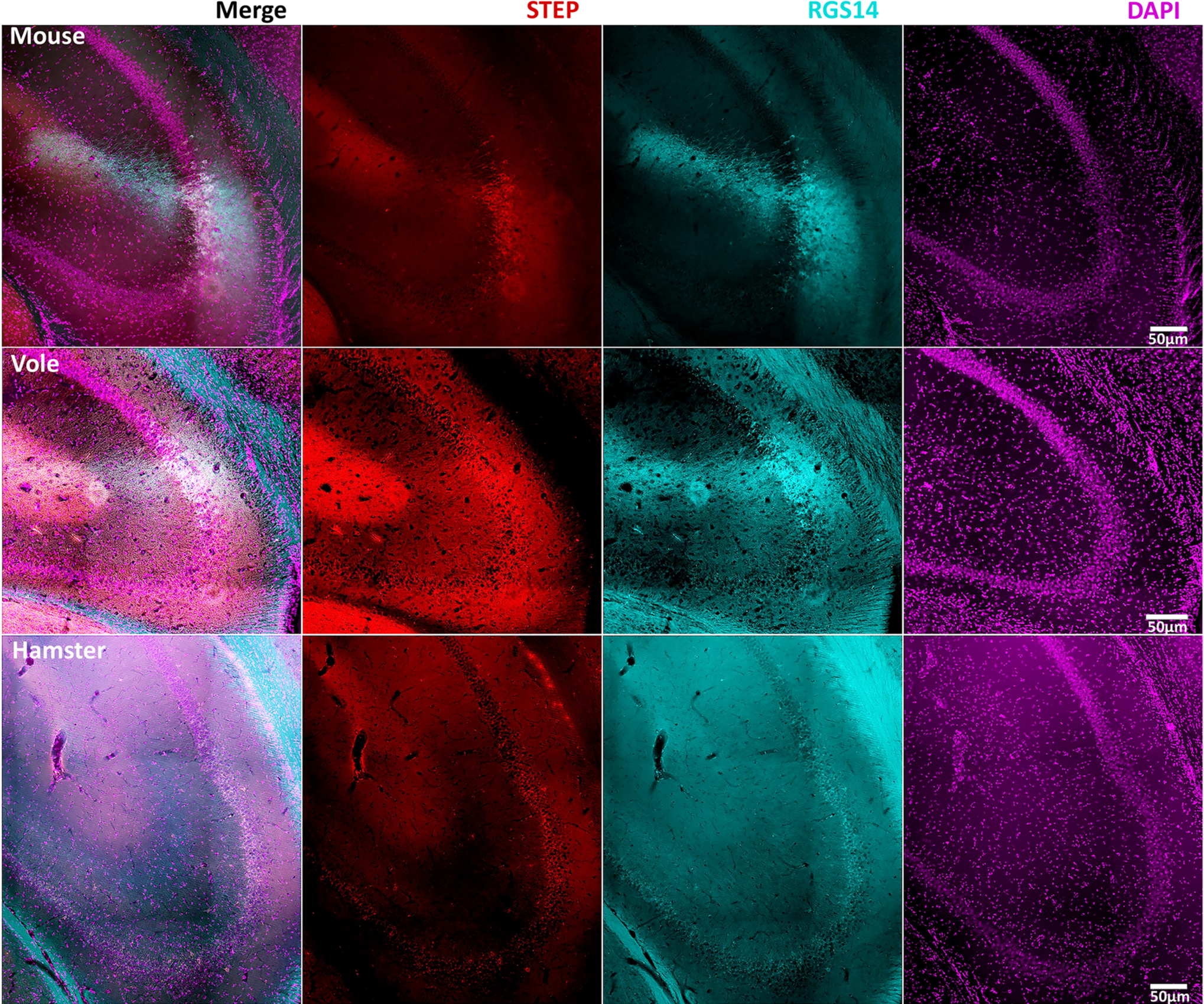
RGS14 colocalizes with a putative area CA2 marker STEP. Striatal-enriched protein tyrosine phosphatase (STEP) is present in dorsal CA2 in all three species and colocalizes with RGS14. STEP (red) is present in all three species, although it is faint in vole and hamster. RGS14 (cyan) demarcates area CA2, and DAPI (magenta) shows cellular density. Partial coronal hippocampal sections are shown.

**Figure 5:**
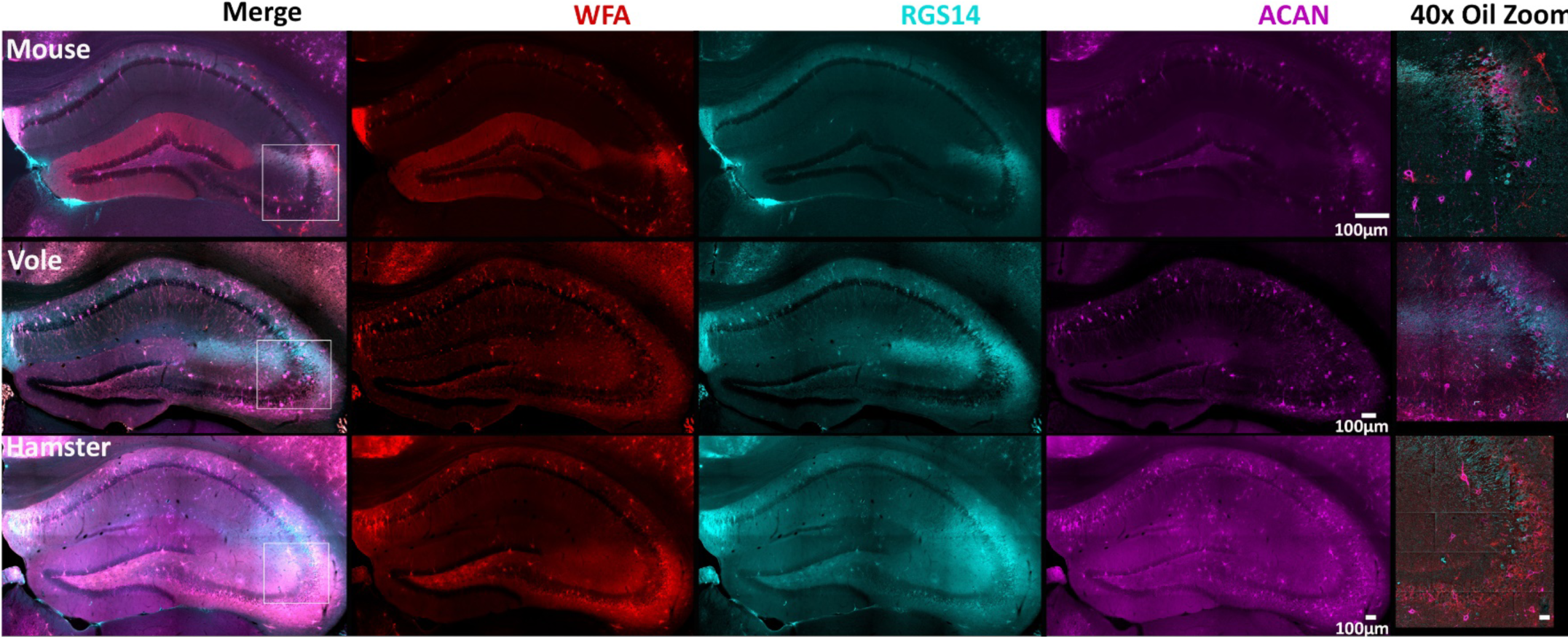
ACAN and WFA stains demonstrate the presence of perineuronal nets on presumptive parvalbumin neurons in all three species and in area CA2 of mouse but not vole or hamster. Staining for *Wisteria floribunda* agglutinin (WFA) and aggrecan (ACAN) is present in all three species and is only concentrated around area CA2 neurons in mouse, but not in vole or hamster. WFA (red) is a N-acetylgalactosamine residue binding lectin that stains neurons in all three species. RGS14 (cyan) indicates area CA2. ACAN (magenta) is one of the main components of perineuronal nets and is indicative of the perineuronal nets. 40x oil zoom-ins of the CA2 and CA3 border are shown in the rightmost column. Coronal hippocampal sections are shown, scale bars on 40x oil images are set to 25μm.

### Both Glucocorticoid Receptors and Mineralocorticoid Receptors are similarly expressed in all three species

The glucocorticoid receptor (GR) and mineralcorticoid receptor (MR) are stress-hormone receptors that, in mouse, are concentrated in CA1 and CA2, respectively (Han et al., 2005; Kalman & Spencer, 2002; McEwen, 1982; Viengchareun et al., 2007). CA2 molecular markers, as well as CA2-dependent synaptic and behavioral phenotypes, are lost in MR knockout mice (Berger et al., 2006; McCann et al., 2021; Ter Horst et al., 2014), demonstrating an integral role of MR in CA2 identity in mice. Therefore, as part of our analysis of whether CA2 exists in hamster and vole, we compared MR and GR expression in hippocampus of all three species.

We stained mouse, vole, and hamster sections with antibodies raised against GR and MR to label neurons expressing these receptors and co-stained tissue with PCP4 to visualize area CA2. In all three species, we observed GR and MR staining in the CA fields (Fig. 6 a-d), with GR enriched in CA1 and MR enriched in CA2. Although the relative distribution of the two receptors between CA1 and CA2 were similar, the overall level of staining for both MR and GR were highest in mouse.

**Figure 6:**
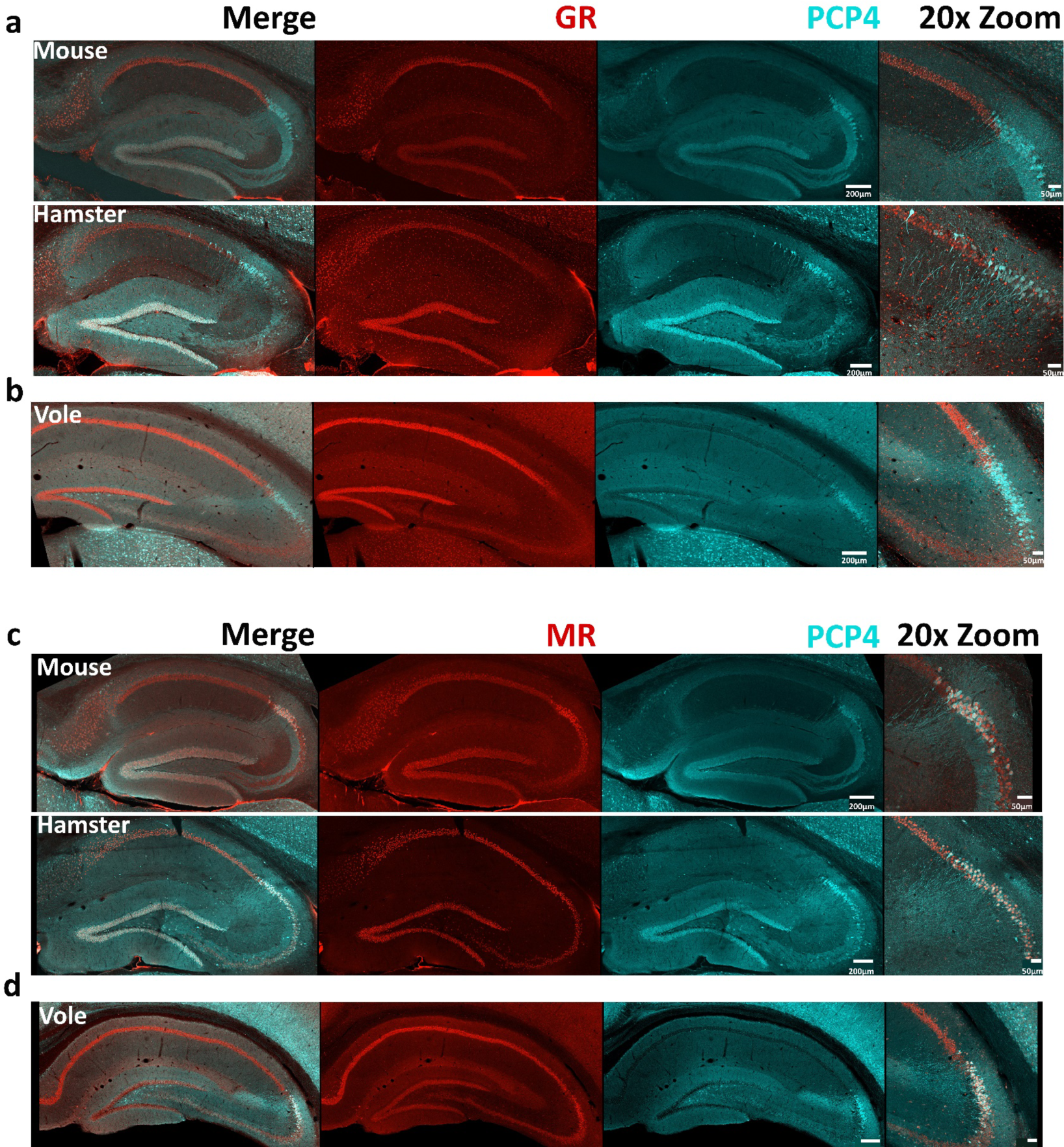
GR and MR are similarly expressed in mouse, hamster, and vole. Staining for GR and MR are present in all three species. a) Whole sagittal hippocampal sections (mouse, hamster) and b) whole coronal hippocampal sections (vole) are shown. GR (red) is present in all three species with little overlap with PCP4 (cyan). c, d) MR (red) is present in all three species, with CA2 neurons (marked by PCP4, cyan) showing some MR enrichment over the other hippocampal regions. Scale bars are as indicated.

## Discussion

This study is the first to characterize hippocampal area CA2 in voles and hamsters, species that are highly useful for studies of social behavior and is important because of the newly-appreciated role of CA2 in rodent social behavior and memory. To understand whether findings in mouse could translate to other species, including those used for prosocial and aggressive behaviors, we first sought to answer the question, “do hamsters and voles have an area CA2?”. Our findings argue that these species do, indeed, have a CA2, or at least a structure that is molecularly and anatomically similar to that in mice. Therefore, a critical role of CA2 in hamster and vole social behaviors is plausible and worthy of further investigation.

We found that, like in mice, prairie voles and Syrian hamsters express RGS14 in a discrete population of hippocampal pyramidal cells that are anatomically positioned between the distal end of the mossy fiber projection from the dentate gyrus and the CA1 pyramidal cells. Thus, we suggest that the area containing these neurons should be considered CA2. Like in mice, RGS14-positive CA2 neurons in voles and hamsters also express PCP4 and STEP, and staining for two receptors found in hippocampus—MR and GR—also presented similar expression patterns amongst the three species. The commonality of these protein expressions across species suggests that many of the characteristics and functions of CA2 neurons may be conserved across species, such as their requirement for social memory and social encoding by place field remapping (Alexander et al., 2016; Samadi et al., 2023; Zhao et al., 2007). However, dissimilar to the case in mouse hippocampus, CA2 pyramidal neurons in voles and hamsters were not unique among the CA subfields in their expression of PNN markers. In fact, WFA and ACAN primarily stained pyramidal neurons in CA3 in these species.

Voles and hamsters are used in studies of social behavior research due to their unique behavioral characteristics. For example, prairie voles exhibit strong pair-bonding behaviors and are specifically aggressive in protection of their mates or pups (Beery et al., 2021; Carter et al., 1995; Gillera et al., 2020). These characteristics make them useful in studies looking at prosocial behaviors. In contrast, male and female hamsters are thought to be solitary in the wild and are naturally antagonistic animals, making them very useful for studies on aggression (Borland et al., 2020; Luckett et al., 2012; Wise, 1974).

Mice serve as model organisms in studies of virtually all areas of biomedical research, including social and non-social behavior, largely owing to the current prevalence of genetic mouse models and the relative ease of genetic manipulation. Thus, a significant amount of our understanding of area CA2’s role in social behavior and memory has come from studies performed in mice. Indeed, CA2 neuronal activity as well as the highly CA2-enriched social neuropeptide receptor vasopressin 1b, specifically, are required for certain types of social memory and aggressive behaviors in mice (Cilz et al., 2019; Hitti & Siegelbaum, 2014; Pagani et al., 2015; Smith et al., 2016; Wersinger et al., 2007; Young et al., 2006). Additionally, MR, which is implicated in some social deficits that are characteristic of neurodevelopmental disorders, is enriched in both mouse and human CA2, and genetic deletion of MR in mice impairs social behavior (Ter Horst et al., 2014; Cukier et al., 2020; McCann et al., 2021).

### Area CA2 in other species

Our finding that vole and hamster have an area CA2 may inform future behavioral studies. Previously used techniques (e.g., Nissl staining), while allowing researchers to visualize cell sizes and density, are limited in that they cannot distinguish cell types based on molecular characteristics. For example, a recent study in foxes (*Vulpes vulpes*) showed staining for PCP4 in a population of neurons that were very likely in CA2, yet no distinct population of CA2 cells were evident with Nissl staining (Dudek et al., 2023). Similarly, studies using Nissl staining, but not IF, have suggested that domestic cats lack a distinct CA2 cell population based on the anatomical definition, where the large pyramidal cells without input from the mossy fibers were spread out and not easily distinguished from the pyramidal cells of area CA3 (Hirama et al., 1997; Zilli et al., 2022). Identification of area CA2 using solely anatomical staining without the additional molecular information that IF provides, has faced similar challenges in domestic dogs, rabbits, guinea pigs and hedgehogs (Hof et al., 1996; Kunzle & Radtke-Schuller, 2001; Potegal et al., 1993; Rami et al., 1987). However, some studies including single-nucleus transcriptomic experiments suggest that dogs do have cells with molecular characteristics like those of cells in area CA2 in other animals (Amayasu et al., 1999; Dudek et al., 2023; Hof et al., 1996; Ragbetli et al., 2010; Zhou et al., 2022). More recent work looking at different species of mole rats have demonstrated PCP4 staining that meets the molecular definitions for area CA2 used here (Stöber & Oosthuizen, 2023). These and future studies of protein expression and anatomical connectivity of CA2 across species will be critical to fully understand how area CA2 contributes to prosocial and aggressive characteristics of animals used in behavioral studies.

### Perineuronal nets in vole and hamster

In mouse hippocampus, pyramidal neurons of CA2, but not CA3 or CA1, are surrounded by PNNs, and several studies suggest PNNs play a role in some social and repetitive behaviors (Briones et al., 2022; Carstens et al., 2021; Balmer et al., 2009; Carstens et al., 2016; Hartig et al., 1992). Consistent with previous work, in mouse we found that PNN staining with both WFA and antibodies against ACAN was strong surrounding mouse CA2 pyramidal cells and some non-pyramidal interneurons throughout the hippocampus (Carstens et al., 2016). However, neither vole nor hamster displayed a concentration of ACAN or WFA stain around CA2 and instead displayed staining primarily in CA3 (with a relatively small portion of vole CA2 being stained), suggesting that PNNs play a species-specific role in regulating behavior that may or may not include social behavior.

PNN expression can dynamically change in response to factors such as social stimuli, environment, and neuronal activity (Dityatev, A, et al., 2007; Carstens et al., 2021; Carstens et al., 2016; Cope et al., 2022; Murthy et al., 2019). Thus, the different social environments experienced by the three species may factor in to the differential PNN distribution, or vice versa. Additionally, neuronal activity levels inversely affect staining for PNNs (Carstens et al., 2021), raising the possibility that activity levels of CA3 and CA2 pyramidal neurons may differ across the three species. Further, PNN expression is thought to reflect the closing of critical periods of plasticity during postnatal development and are thought to curb synaptic plasticity in adulthood in CA2 and elsewhere (Carstens et al., 2021; Noguchi et al., 2017; Fawcett et al., 2019; Pizzorusso et al., 2002). During the first postnatal week in mice, before PNNs appear, LTP can be induced in CA2 neurons, but once PNNs are expressed in the second postnatal week and beyond, LTP is no longer seen in CA2 (Carstens et al., 2016; Cope et al., 2022; Noguchi et al., 2017). However, enzymatic degradation of PNNs after the second postnatal week permits LTP in CA2 (Carstens et al., 2016). Therefore, PNNs appear to limit synaptic plasticity in mouse CA2 and may similarly limit synaptic plasticity in vole and hamster CA3. That said, PNNs are only one component of the complex LTP-regulating machinery in CA2; RGS14, PCP4 and STEP also regulate LTP and begin to be expressed postnatally around the same ages in mice (Carstens & Dudek, 2019; Laham et al., 2021). Although whether CA3 and CA2 exhibit LTP in hamster and vole is unknown, our findings present a curious possibility of varying propensities for LTP in CA3 and CA2 and a differential role of PNNs in social behavior across the three species.

### Molecular markers of area CA2 and other hippocampal regions

RGS14, which acts as an endogenous regulator of synaptic plasticity in CA2 (Lee, et al., 2010, Evans et al., 2018), demarcates most clearly the CA1-CA2 border and, based on our results, also appears to be an excellent CA2 marker in voles and hamsters. Some studies performed on tissue from human and non-human primates have also found enriched RGS14 staining in CA2 (Carstens et al., 2021; Squires et al., 2018), future strengthening the argument that antibodies against RGS14 can be used as a CA2 marker in a variety of species. Similarly, staining for PCP4 delineates well the CA2-CA3 border well in the species studied here, and both PCP4 and STEP colocalize well with RGS14 (Radzicki et al., 2023).

ZnT-3, which is necessary for the accumulation of zinc ions inside synaptic vesicles, is a useful marker of the axons from dentate granule cells (i.e., mossy fibers) that sweep across CA3 and terminate in CA2 (Kohara et al., 2014; Palmiter et al., 1996; Sindreu et al., 2011; Sindreu & Storm, 2011). Our results show that there are some differences in mossy fiber staining with ZnT-3 amongst species, with ZnT-3 strongly staining mossy fibers in mouse and weakly staining mossy fibers in vole and hamster. This difference may reflect truly different expression levels, possibly with different functional effects in the hippocampal circuitry, but more likely, the different staining intensity reflects differences in the antibody binding across species in that many of the antibodies used in this study were monoclonal antibodies. In contrast to polyclonal antibodies, monoclonal antibodies are typically targeted against only one protein epitope, which could differ substantially between different species and thus differentially affect binding.

A polyclonal antibody against calbindin, however, robustly stained mossy fibers in all three species (Inoue et al., 1998; Takeda et al., 2001). Calbindin, which functions as a sensor and transporter of intracellular calcium, shows pyramidal neurons in CA1, mossy fibers in CA3, and granule cells in dentate (Mody et al., 1987; Rami et al., 1987). Calbindin is also differentially expressed in rodents and primates; it is highly expressed in mouse CA1 but not human CA1, and, interestingly, some calbindin-positive pyramidal neurons have been found in primate CA2 (Ding & Van Hoesen, 2015; Merino-Serrais et al., 2020). Here, we found CA1 neurons in mouse and hamster stained for calbindin, but CA1 neurons were not stained in vole despite the mossy fibers staining for calbindin. This result was similarly reported in guinea pig (Rami et al., 1987). Thus, vole is not unique in its lacking a double layer of calbindin positive and negative cells in CA1 (Rami et al., 1987).

GRs and MRs bind the stress hormones cortisol (in human) and corticosterone (in rodents) and act as transcription factors that control a variety of downstream transcriptional networks when activated (McCann et al., 2021; McEwen, 1992; Viengchareun, et al., 2007). GRs and MRs expressed in the brain mediate the behavioral and physiological response to stress, emotional states, and learning and memory. In mouse and human hippocampus, the highest ratio of MR to GR expression is found in area CA2, making staining for MRs and GRs acceptable markers of CA2 neurons and the CA1/CA2 border, respectively (Herman et al., 1989; Han et al., 2005; Kalman & Spencer, 2002; McCann et al., 2021). MRs are also key in determining the overall molecular profile of CA2. In mice, genetic knockout of MRs results in loss of most other CA2 markers in the region and upregulation of CA1 markers such as GR and WFS1 (McCann et al., 2021). Given the rich CA2 expression of MRs in hamster and vole, CA2 marker expression in these species may similarly be under the transcriptional control of MR and may point to a role of CA2 in stress-related behaviors across multiple species.

### Implications for CA2’s role in social behavior

CA2 plays a critical role in social behavior. In terms of protein expression, CA2 expresses an abundance of vasopressin 1b and oxytocin social neuropeptide receptors as well as the mineralocorticoid stress hormone receptor (Lin et al., 2018; McCann et al., 2021; Tirko et al., 2018; Wersinger et al., 2002; Young et al., 2006). CA2 pyramidal neurons also express ACAN, which yields PNNs, with a developmental onset of expression paralleling a developmental closure of the critical period of synaptic plasticity and a shift of pup’s preference for a maternal caregiver to a novel conspecific (Cope et al., 2022; Laham et al., 2021). In adult animals, CA2 neuronal activity is modified by exposure to a social stimulus and is required for social learning, memory, and aggression (Alexander et al., 2018; Alexander et al., 2016; Hitti & Siegelbaum, 2014; Leroy et al., 2018; Pagani et al., 2015). In addition, knocking out the vasopressin 1b receptor reduces aggressive behavior and impairs social recognition in male mice, and the conditional deletion of oxytocin receptors in both CA2 and CA3 impairs long-term social recognition in mice (Lin et al., 2018; Pagani et al., 2015; Wersinger et al., 2002).The current body of work supports the similarity of CA2’s molecular profile and potential function in multiple species. However, the intriguing difference in PNN expression across the three species speaks to potential insights into the differences in social behavior.

MRs have been shown to be a key factor in determining CA2’s molecular profile and has been implicated in social deficits, such as social recognition memory, which is impaired in MR knockout mice (Berger et al., 2006; McCann et al., 2021; Ter Horst et al., 2014). Furthermore, mutations in *NR3C2*, the gene that encodes MR, are associated with autism spectrum disorder in humans, further strengthening the implications that MR dysregulation, and perhaps CA2, may be involved in some the behavioral deficits found in certain neurodevelopmental disorders (Cukier et al., 2020; McCann et al., 2021).

Being able to form, assess, and remember social connections and interactions with conspecifics is key to the survival of many species, including ours. These skills are disrupted in neurodevelopmental disorders such as autism and schizophrenia (Barendse et al., 2018; Barkl et al., 2014; Modi et al., 2019) and is impaired in neurocognitive and neurodegenerative disorders such as some forms of dementia (Churchyard & Lees, 1997; Dickson et al., 1991; Kalaitzakis et al., 2009). Likewise, one of the leading causes of inpatient care for a subset of individuals with autism is aggressive behavior, including caregiver aggression and self-injury (such as hitting, biting, and scratching), especially after overstimulation or increased stress (Brown et al., 2023; Hirota et al., 2020; Im, 2021; Pedersen et al., 2018). Because of the role CA2 plays in aggression, social learning, and memory, multiple labs are now investigating whether CA2 is implicated in disorders with an impaired social component or increased risk of aggressive tendencies (Carstens et al., 2021; Modi et al., 2019; Yang et al., 2021). Knowing the position and molecular make-up of CA2 in both vole and hamster, which are useful animal models used in studies of prosocial behaviors and aggression, can provide researchers with key information in the pursuit of addressing how social behaviors are controlled in these animals.

## Conclusions

In this study, we used IF and commonly used antibodies raised against proteins highly expressed in mouse hippocampal area CA2 to investigate whether voles and hamsters had cellular populations analogous to mouse CA2. We identified RGS14 as a reliable marker of CA2 in the three species. As in mice, RGS14-positive cell populations were found at the end of the mossy fiber projections (most robustly visualized with calbindin) in voles and hamsters. This cellular population also colocalized well with two other common CA2 markers, PCP4 and STEP. Staining for receptors commonly found in mouse hippocampus, GR and MR, also showed similar expression amongst the three species with GR expression highest in CA1 and MR enrichment in CA2 of all three species. Staining for ACAN and WFA was present in all three species but did not concentrate around CA2 pyramidal cells in vole or hamster. Rather, we observed PNN staining primarily in CA3 in tissue from those animals. The many similarities in CA2-specific protein expression across species argues for the presence of a CA2 in voles and in hamsters and provides a platform for understanding the species-specific roles of CA2 in social behaviors.

## Acknowledgements

This research was funded by the Intramural Research Program of the National Institute of Environmental Health Sciences, U.S. National Institutes of Health (ES 100221), by MH062044 and MH122622 to KLH, and by AR160055 to HBP. We thank the expert staff the NIEHS Fluorescence Microscopy and Imaging Center for their help with imaging and Dr. Meilan Zhao for performing some of the immunofluorescence staining. We also thank members of the Dudek, Patisaul, and Huhman labs for their careful reading of the manuscript.

## Notes

**Conflict of Interest Statement:** The authors have no conflicts to declare.

### Competing Interest Statement

The authors have declared no competing interest.

